# SPaRTAN, a computational framework for linking cell-surface receptors to transcriptional regulators

**DOI:** 10.1101/2020.12.22.423961

**Authors:** Xiaojun Ma, Ashwin Somasundaram, Zengbiao Qi, Harinder Singh, Hatice Ulku Osmanbeyoglu

## Abstract

The identity and functions of specialized cell types are dependent on the complex interplay between signaling and transcriptional networks. We present SPaRTAN (Single-cell Proteomic and RNA based Transcription factor Activity Network), a computational method to link cell-surface receptors to transcription factors (TFs) by exploiting cellular indexing of transcriptomes and epitopes by sequencing (CITE-seq) datasets with cis-regulatory information. SPaRTAN is applied to immune cell types in the blood to predict the coupling of signaling receptors with cell context-specific TFs. The predictions are validated by prior knowledge and flow cytometry analyses. SPaRTAN is then used to predict the signaling coupled TF states of tumor infiltrating CD8^+^ T cells in malignant peritoneal and pleural mesotheliomas. SPaRTAN greatly enhances the utility of CITE-seq datasets to uncover TF and cell-surface receptor relationships in diverse cellular states.

**Significance:** Recently single-cell technologies such as CITE-seq have been developed that enable simultaneous quantitative analysis of cell-surface receptor expression with transcriptional states. To date, these datasets have not been used to systematically develop cell-context-specific maps of the interface between signaling and transcriptional regulators orchestrating cellular identity and function. We developed a computational framework, SPaRTAN (Single-cell Proteomic and RNA based Transcription factor Activity Network) that integrates single-cell proteomic and transcriptomic data based on CITE-seq with cis-regulatory information. We applied our method to publicly available (PBMCs) and new CITE-seq datasets (mesothelioma). Our predicted TF activity and cell-surface receptor relationships are validated by prior knowledge as well as experimental testing. SPaRTAN reveals many previously unidentified surface-receptor associated TFs.

## Introduction

The reciprocal interplay between complex signaling inputs and transcriptional responses dictate the generation of distinct cell types and their specialized functions. Dysregulation of this interplay leads to the development and progression of disease, most clearly delineated in the context of certain cancers, chronic infections and autoimmune diseases. Understanding these dynamic programs at the single-cell level represents a formidable challenge. Emerging single-cell genomic technologies (1-4) provide a transformative platform to characterize, in a comprehensive and unbiased manner, the full range of cell types and their genomic programming in health and disease.

The computational prediction of gene regulatory programs based on single-cell genomic datasets is a relatively new field. There is still a large methodological gap between generating single-cell datasets and delineating cell-specific regulatory programs orchestrating cellular identity and function. Early gene regulatory program inference methods use single-cell RNA-seq (scRNA-seq) data alone or in combination with TF motifs in annotated promoter regions (5-9). These methods primarily depend on co-expression of TFs and their potential target genes (6) and thus are not suitable for many TFs whose transcripts are expressed at low levels or whose activities are post-transcriptionally regulated. Moreover, co-expression may not always imply co-regulation. With the recent availability of single-cell epigenomic datasets, the tools of regulatory genomics are being applied to infer TFs associated with accessible chromatin regions (10) at both promoter-proximal as well as distal regions and in turn with gene expression (11). However, these approaches do not comprehensively consider the relationships between signaling systems (e.g., from proteomic data) and transcriptional states of individual cells. Recent breakthroughs in single-cell genomics have linked single-cell gene expression data with quantitative protein measurements using index sorting (12) and barcoded antibodies (1, 2), in particular *cellular indexing of transcriptomes and epitopes by sequencing* (CITE-seq) (1). CITE-seq adds a step in which barcoded antibodies – a second set of barcodes – are incubated with the single-cell suspension into droplet based scRNA-seq protocol. These barcodes, which have polyA tails, are then linked to the barcodes from beads at the same time mRNAs are linked. Reads containing barcodes associated with each bead are separated by cell. Then, reads that align to transcripts are used to quantify mRNA levels while those from barcoded antibodies (antibody-derived tags (ADTs)) are used to quantify protein levels. To date, the datasets generated by this powerful platform have not been used to link the expression of cell surface proteins, e.g., signaling receptors, with the activities of TFs and gene expression programs in individual cells.

Here, we describe a computational framework for exploiting single-cell proteomic (scADT-seq) and corresponding single-cell transcriptomic (scRNA-seq) datasets, both obtained using CITE-seq, to link expression of surface proteins with inferred TF activities. Our framework, SPaRTAN (**S**ingle-cell **P**roteomic **a**nd **R**NA based **Tr**anscription factor **A**ctivity **N**etwork), advances our prior algorithmic approach based on bulk tumor datasets (13-15). SPaRTAN model views expression of surface proteins (ADT counts) as a proxy of their activities; signaling emanating from these proteins converges on particular TFs, whose activities, in turn, regulate the expression of their target genes (**Fig. 1**). More specifically, we use a regularized bilinear regression algorithm called affinity regression (AR) (16) to learn an interaction matrix between upstream cell-surface receptor proteins and downstream TFs that predicts target gene expression. The trained SPaRTAN model can then infer the TF activity given a cell’s surface protein expression profile or infer the cell-surface receptor expression given a cell’s gene expression profile. We apply and experimentally test SPaRTAN using CITE-seq datasets from peripheral blood mononuclear cells (PBMCs) and then illustrate its broader utility by predicting signaling coupled TF activities in tumor infiltrating CD8^+^ T cells in the context of malignant peritoneal and pleural mesothelioma.

**Fig. 1:**
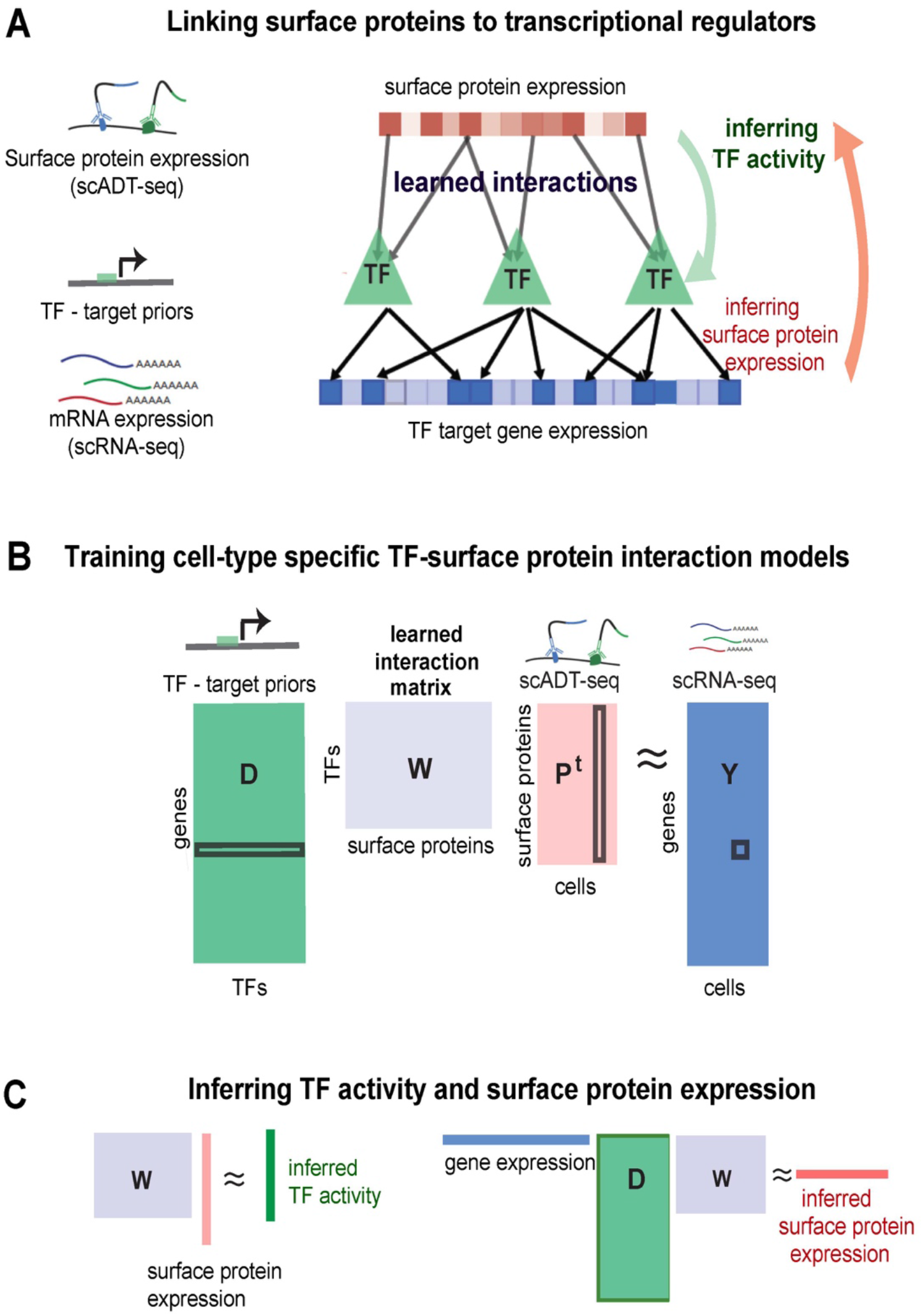
Integrative computational model linking cell-surface receptors to transcriptional regulators -application to peripheral blood mononuclear cells (PBMC) **(A)** Our integrative model (SPaRTAN, Single-cell Proteomic and RNA based Transcription factor Activity Network) utilizes single-cell multi-omics data from cellular indexing of transcriptomes and epitopes by sequencing (CITE-seq) datasets and infers the flow of information from cell-surface receptors to transcription factors (TFs) to target genes by learning interactions between cell surface receptors and TFs that best predict target gene expression. **(B)** SPaRTAN trains on mRNA and surface protein expression data from a set of cells, along with curated TF target-gene interactions, to learn a model that links upstream signaling to downstream transcriptional responses. Specifically, the algorithm learns a weight matrix **W** between cell-surface proteins and TFs that predicts gene expression (**Y**) from cell-specific surface protein levels (**P**) and TF target-gene interactions (**D**) by solving the bilinear regression problem shown. **(C)** Using the trained interaction matrix, one can predict TF activities from the cell surface protein expression profile, or protein activities from the cellular mRNA expression data and the TF-target gene hit matrix.

## Results

### SPaRTAN learns a cell-type specific interaction model for cell-surface receptors and TFs

SPaRTAN integrates parallel single-cell proteomic and transcriptomic data (based on CITE-seq) with cis-regulatory information (e.g., TF:target-gene priors) for predicting cell-specific TF activities and surface protein expression for linking surface receptor signaling to downstream TFs. Formally, we used an affinity regression (AR) (16) algorithm, a general statistical framework for any problem where the observed data can be explained as interactions between two kinds of inputs, to establish an interaction matrix (**W**) between surface reeceptors/proteins (**P**) and TFs (**D**) that predicts target gene expression (**Y**) (**Fig. 1B**). To determine the set of TFs that potentially regulate each gene (**D**), we utilized curated TF target-gene interactions (17). We trained independent SPaRTAN models for each cell type that explain gene expression across cells (**Y**) in terms of surface protein expression (**P**) and TF target-gene interactions (**D**) (see **Methods** for details).

We can use the trained interaction matrix (**W**) to obtain different views of a CITE-seq data set; *e*.*g*., to predict TF activity from a cell’s surface protein expression profile (**WP**^T^) or to predict surface protein expression from a cell’s gene expression profile (**Y**^T^**DW**) (**Fig. 1C**). Intuitively, information flows down from observed surface protein levels through the learned interaction matrix to infer TF activities and observed mRNA expression levels or propagates up through the TF target-gene edges and interaction network to infer surface protein expression. Importantly, we can use these predicted TF activities and surface protein expression to gain biological insights into different cell types and states as described below.

To evaluate our approach, we first trained cell-type specific SPaRTAN models using an existing peripheral blood mononuclear cells (PBMCs) CITE-seq dataset (**Supplemental Table 1**). Cell types were identified using both protein and gene expression data, the latter with marker genes (see **Methods section, Supplementary Fig. 1-2**). For statistical evaluation, we computed the mean Spearman correlation between predicted and measured gene expression profiles on held-out samples using 5-fold cross-validation for each cell-type specific SPaRTAN model using equal numbers of cells. We obtained significantly better performance than a nearest-neighbor approach based on Euclidean distance in the surface protein profiles (input space) (*P*<0.001, one-sided Wilcoxon’s signed-rank test) as shown in **Fig. 2A**. We next evaluated the approach using an independently generated PBMC CITE-seq dataset (see validation PBMC dataset, **Supplemental Table 1**) and attained similar performance results (**Supplementary Fig. 3**). We also used the PBMC-trained SPaRTAN cell-type specific models, to infer surface protein expression (**Y**^T^**DW**) for each cell type in training and validation CITE-seq PBMC data sets (see **Supplemental Table 1**). For surface proteins whose Spearman correlations between measured and inferred activities were above 0.3 on training dataset, we found similarly strong correlations between measured and predicted surface protein levels on the validation CITE-seq data (**Supplementary Fig. 4**).

**Fig. 2:**
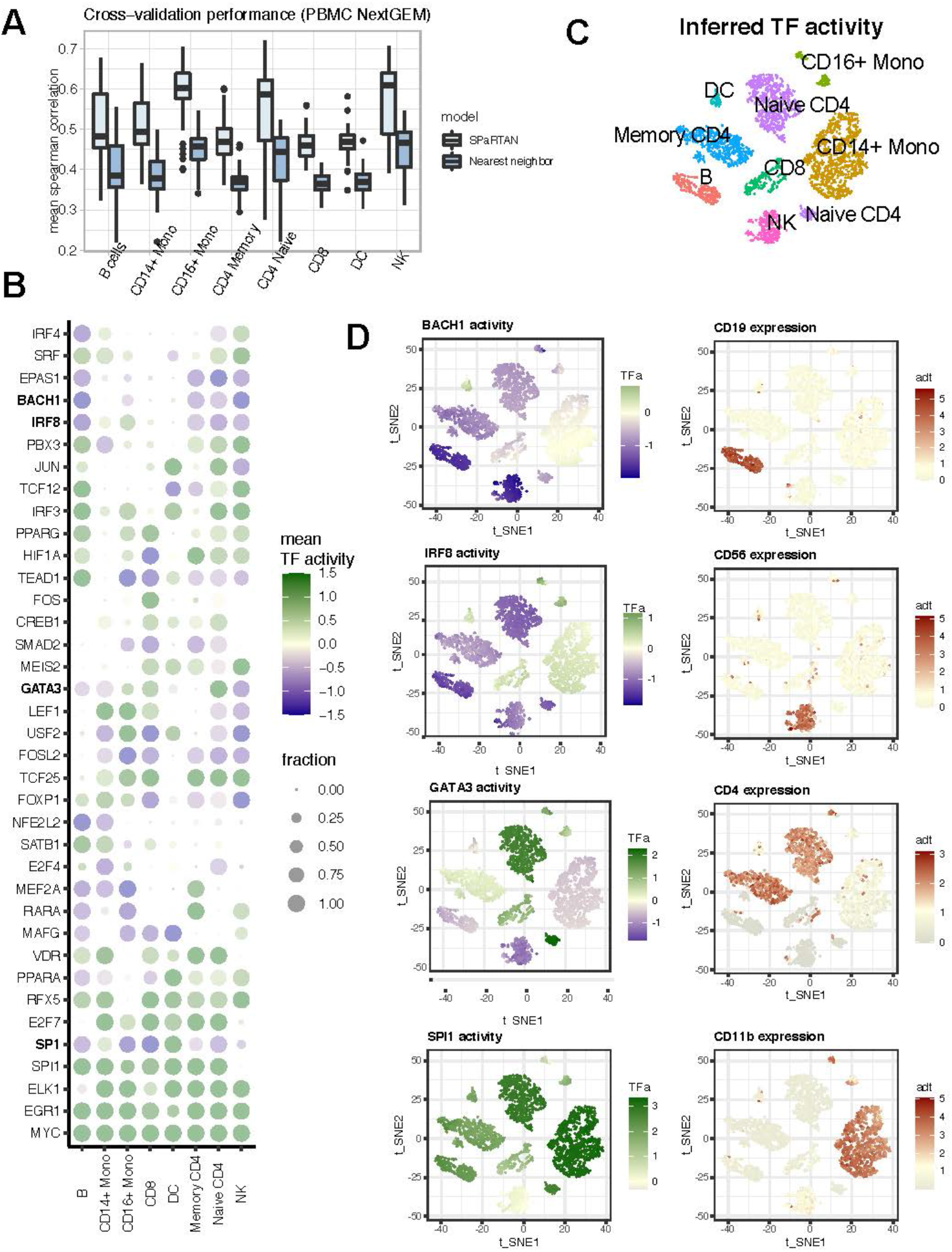
SPaRTAN identifies cell-type specific TFs in peripheral blood mononuclear cells (PBMC) **(A)** SPaRTAN accurately predicts relative gene expression on held-out PBMC (10X Genomics, NEXTGEM, training dataset) cells for each cell-type. Performance of the SPaRTAN models for each PBMC cell type compared to nearest neighbor methods. Boxplots showing mean Spearman correlations between predicted and actual gene expression using the SPaRTAN model (light blue); nearest neighbor by surface protein expression profile (blue) (*y*-axis) for PBMC CITE-seq data from 10X Genomics (Next GEM) each cell-type. **(B)** Dot plot showing the median TF activity z-score of TFs across different cell types. The dot size indicates a fraction of cells for which indicated TF is identified as a significant regulator within designated cell type. For clarity, the union of the top 8 most prevalent significant TFs in each cell type-specific model is shown. **(C)** t-SNE on the inferred TF activity matrix. Cells are colored according to major cell types. **(D)** BACH1, IRF8, GATA3, and SPI1 inferred TF activity and CD19, CD56, CD4 and CD11b protein expression overlay on t-SNE of TF activities.

### SPaRTAN identifies cell type-specific TFs

Next, we used our approach to predict cell-type specific activities of TFs (**WP**^**T**^) in PBMCs. To assess the statistical significance of inferred TF activities, we developed an empirical null model based on randomly permuted gene expression profiles for each cell-type (see **Methods section**). Then, we asked whether the value of individual TF activities for each cell were significantly low or high relative to the corresponding distribution over permuted data. We corrected for multiple hypotheses across TFs and identified significant shared and cell-type specific TFs. **Fig. 2C** shows the fraction of cells per cell-type where each TF was identified as a significant regulator; the representation encompasses the top 8 most prevalent significant TFs for each indicated cell type. **Fig. 2C-D** show the inferred activity distribution of four TFs identified from our analysis: *BACH1, GATA3, SPI1*, and *IRF8* with overlay of CD19, CD56, CD4 and CD11b surface protein expression. These inferred TFs are known regulators of several of the cell types within the PBMCs including *GATA3* (18) for naïve CD4^+^ and CD8^+^ T cells; *SPI1* (*PU1*) (19, 20) for B, CD14^+^/CD16^+^ monocytes and dendritic cells; *BACH1* (21) for B, NK, and CD 4 T cells; *PRDM1* (*BLIMP1*) (22) for B, CD16^+^ monocytes and T cells. TFs involved in cellular activation and proliferation like *EGR1* and *MYC* were shared across all cell types (**Fig. 2D**). Importantly, in spite of accurately inferring their activities we did not reliably detect mRNAs for many cell-type-specific TFs, given their low levels of transcript expression (**Supplementary Fig. 5**).

### SPaRTAN delineates cell type-specific TFs coupled with cell-surface receptors

To explore the associations between inferred TF activities and surface protein expression at a single-cell level, we first computed Pearson correlation coefficients (PCC) between (inferred) TF activity and surface protein expression for each TF-surface protein pair within each cell-type. **Fig. 3A-C** (see also **Supplemental Fig. 6**) shows the two-way clustering of TFs and proteins by these pairwise PCC in B, CD8^+^ and CD4^+^ T memory cells. We identified several known as well as novel TF-surface protein relationships for each cell type. We next evaluated the known pathway overlap between TF and surface proteins. Correlated pairs were enriched for known pathways for most cell types (*P* < 10^−3^ by the hypergeometric test, minimum one overlapping pathways) compared to the alternative approach(6) for identification of cell-type specific TF activities using only single-cell gene expression measurement (**Supplemental Table 2)**. Importantly, the analysis suggests that a cell-surface receptors can couple with shared and context-dependent downstream transcriptional regulators in different cell types (**Fig. 3D-F**). For example, SPaRTAN-predicted *FOSL2* activity was correlated with CD27 (member of the TNF receptor super family (23)) protein expression in B, CD8^+^ and CD4^+^ memory T cells. Recent studies have shown that *FOSL2* represses Treg development and controls autoimmunity (24) and can also control autoreactive B cells in patients with Systemic lupus erythematosus (SLE) (25). In contrast, SPaRTAN-predicted *STAT5A* activity was highly correlated with CD27 protein expression in B and CD4^+^ memory T cells and with TIGIT (inhibitory immunoreceptor targeted in antitumor immunotherapy (26)) protein expression in CD8^+^ T cells.

**Fig. 3:**
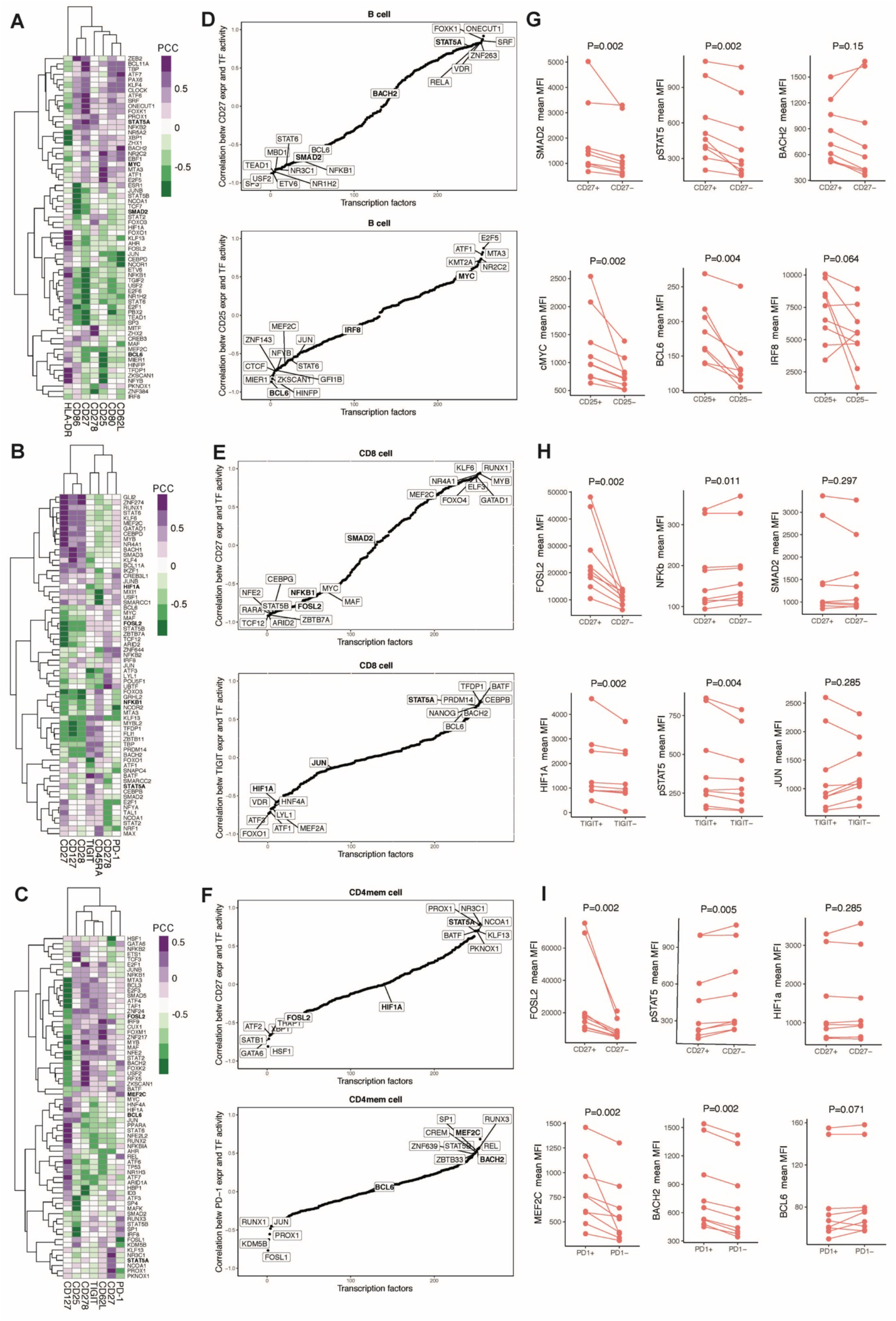
Analysis of SPaRTAN inferred TF activities with cell surface receptor expression in PBMCs - validation with multiparameter flow cytometry. Heatmap revealing correlations between inferred TF activities (rows) and surface protein expression (columns) in **(A)** B cells, **(B)** CD8^+^ T cells, and **(C)** CD4^+^ memory T cells. For clarity, surface proteins with Pearson’s correlation coefficient (PCC) values with TFs below 0.75 are filtered, and then the union of the top 10 most correlated TFs with each surface protein is shown for each cell type. Representative sorted correlation plots between **(D)** CD27 (top) CD25 (bottom) protein expression and inferred TF activities across B cells; **(E)** CD27 (top) TIGIT (bottom) protein expression and inferred TF activities across CD8^+^ T cells; **(F)** CD27 (top) PD-1 (bottom) protein expression and inferred TF activities across CD4^+^ memory T cells. **(G-I)** Validation of predictions using flow cytometry analysis. Briefly, B, CD8^+^ and CD4^+^ memory T cells were isolated from peripheral blood (PBL) from healthy donors (n=9) and stained for indicated surface receptors and intracellular TFs. Paired analysis of TF expression was assessed on surface-protein^+^ and surface-protein^-^ cells. P-values are calculated using paired Wilcoxon signed rank test. Representative validation results are shown in each case with two TFs showing high correlation and a control TF showing low correlation.

To directly test SPaRTAN predicted relationships between cell surface signaling proteins and TF activities in specific cell types, we performed flow cytometry analysis for select surface receptor-intracellular TF pairs using PBMCs from healthy donors (**Fig. 3G-I, Supplementary Fig. 7)**. Consistent with our predictions and in spite of the variation among individual donors, we observed increased expression of *FOSL2* in CD27^+^ CD8^+^ T and CD4^+^ memory T cell subsets compared with their CD27^−^ counterparts; increased expression of *STAT5* in CD25^+^ (IL-2 receptor) B cells compared with the CD25^−^ subset and increased expression of *MEF2C* and *BACH2* with the expression of PD-1 (inhibitory immunoreceptor (27)) in CD4^+^ memory T cells. Importantly, these experimentally validated relationships were not predicted by the alternative approach (6) for identification of cell-type specific TF activities using only single-cell gene expression measurements (**Supplementary Fig. 8)** as well as using only TF mRNA levels (**Supplementary Fig. 9)**.

### SPaRTAN identifies cell state specific TFs coupled with cell-surface receptors

Next, we asked whether our method could be used to analyze distinct cellular states within a given cell type. **Fig. 4A** shows the clustering of cells by cell surface protein expression (excluding cell lineage marker surface proteins), together with inferred TF activities for the same cell ordering, as derived from the CD8^+^ T cell model. Unsupervised clustering by the surface protein (ADT) expression profiles identified five major cellular states of CD8^+^ T cells. In particular, Cluster 1 was distinguished by high expression for CD45RA and low expression for CD45RO which is characteristic of naïve CD8^+^ T cells. Among the most significant differences in inferred TF activity associated with this cluster were *KLF13, FOSL1, USF2, FOXO3, ZEB1, and SMAD5* (FDR-corrected *p* < 10^−5^, t-test; **Fig. 4B**). Cluster 2 was distinguished by cells with high expression of CD27 and CD127 which are characteristic of memory CD8^+^ T cells (28, 29). Among the most significant differences in inferred TF activity associated with this cluster were *RUNX1, SMAD3*, and *BCL11A* (FDR-corrected *p* < 10^−5^, t-test, **Fig. 4B**).

**Fig. 4:**
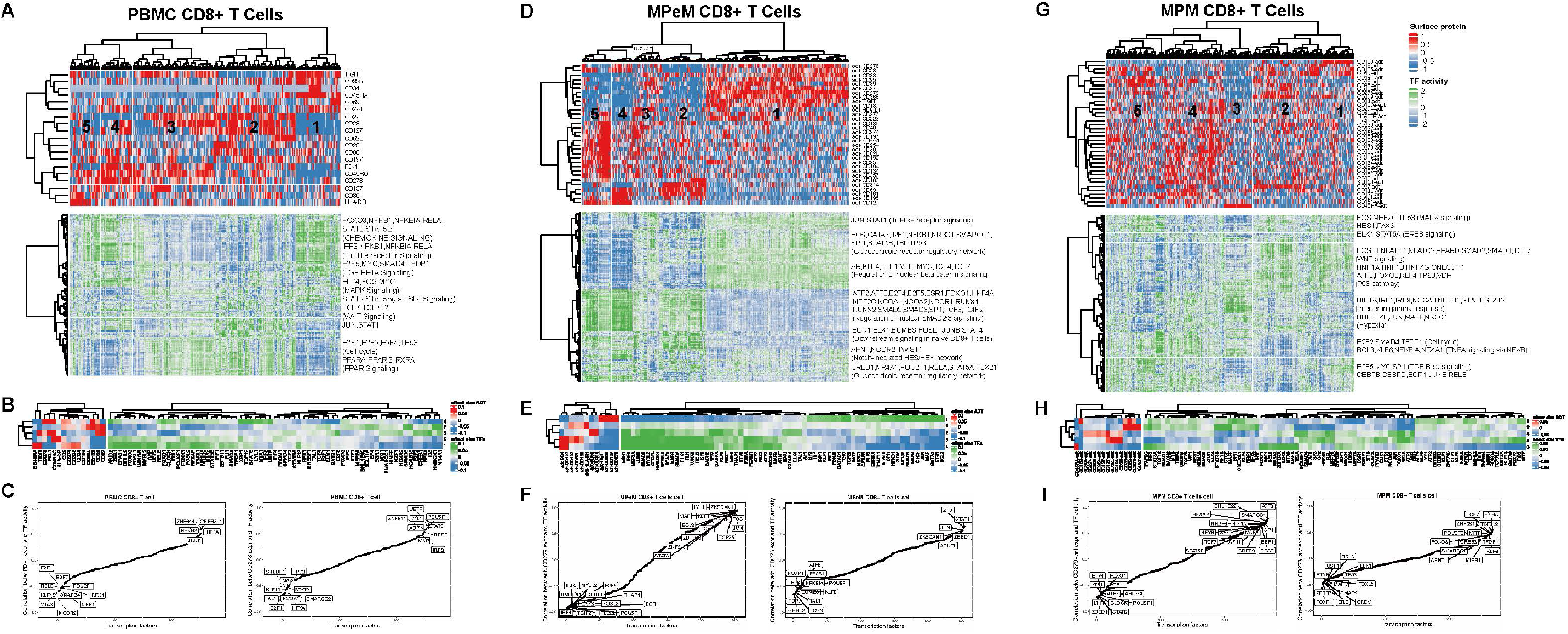
SPaRTAN modeling and analysis of regulatory states of circulating and tumor infiltrating CD8 T cells. **(A)** SPaRTAN model trained on 319 CD8^+^ T cells from the 10X Genomics PBMC dataset. The top heat map shows cells clustered by the surface protein expression (excluding cell lineage maker surface proteins) The bottom panel shows inferred TF activities for each cell based on clustering by surface protein expression. **(B)** Heatmap shows the mean surface protein expression and inferred TF activity between cells in a given cluster vs. those in all other clusters. For each comparison, the absolute value of the mean surface protein expression and inferred TF activity (effect sizes) are ranked and the union of the top 20 TFs for each comparison is shown in the heatmap. **(C)** Sorted correlation plots between PD1 (left) CD278 (ICOS) (right) protein expression and inferred TF activities across PBMC CD8^+^ T cells. **(D)** We trained a SPaRTAN model on MPeM CD8^+^ T cells. The top heat map shows cells clustered by the surface protein expression (excluding cell lineage maker surface proteins) The bottom panel shows TF activities for each cell based on clustering by surface protein expression. **(E)** Heatmap shows the mean surface protein expression and inferred TF activity between cells in a given cluster vs. those in all other clusters. For each comparison, the absolute value of the mean surface protein expression and inferred TF activity (effect sizes) are ranked and the union of the top 20 TFs for each comparison is shown in the heatmap. **(F)** Sorted correlation plots between CD279 (PD1) (left) CD278 (ICOS) (right) protein expression and inferred TF activities across MPeM CD8^+^ T cells. **(G)** SPaRTAN model trained on MPM CD8^+^ T cells. The top heat map shows cells clustered by the surface protein expression (excluding cell lineage maker surface proteins) The bottom panel shows TF activities for each cell based on clustering by surface protein expression. **(H)** Heatmap shows the mean surface protein expression and inferred TF activity between cells in a given cluster vs. those in all other clusters. For each comparison, the absolute value of the mean surface protein expression and inferred TF activity (effect sizes) are ranked and the union of the top 20 TFs for each comparison is shown in the heatmap. **(I)** Sorted correlation plots between CD279 (PD1) (left) CD278 (ICOS) (right) protein expression and inferred TF activities across MPM CD8^+^ T cells.

We performed similar analyses for other cell types (**Supplementary Fig. 10–15**). For example, unsupervised clustering by surface protein expression identified five major cellular states within the B cell compartment (**Supplementary Fig. 10)**. In particular, Cluster 2 was distinguished by B cells with high expression of CD80 which is characteristic of activated B cells (30). Among the most significant differences in inferred TF activity associated with this cluster were *ARID3A, TFDP1* and *TP63* (FDR-corrected *p* < 10^−5^, t-test; (**Supplementary Fig. 10B**). Cluster 3 was distinguished by B cells with high expression of CD34 and low expression of CD27 and CD80 which are characteristics of naïve B cells. Among the most significant differences in inferred TF activity associated with this cluster were *TEAD1, RFX5, JUNB, PBX2, FOXO3*, and *HMBOX1* (FDR-corrected *p* < 10^−5^, t-test (**Supplementary Fig. 10B**). Moreover, Cluster 4 was distinguished by B cells with high expression of CD28 and CD27 and low expression of CD20 which are characteristics of plasmablasts. Among the most significant differences in inferred TF activity associated with this cluster were *MITF, STAT5A*, and *FOS* (31-33) (FDR-corrected *p* < 10^−5^, t-test, **Fig**. (**Supplementary Fig. 10B**).

To illustrate the broader utility of our approach, as a discovery platform, we applied it to the tumor microenvironments (TMEs) of malignant peritoneal (MPeM) and pleural (MPM) mesothelioma. CITE-seq data sets for cells within these TMEs were generated (**Supplementary Fig. 16-17**) and queried for the regulatory states of tumor infiltrating CD8^+^ T cells. Unsupervised clustering using cell surface protein expression patterns identified five major populations of MPeM CD8^+^ T cells (**Fig. 3D**). We tested for statistical differences in inferred TF activities and surface protein expression in a given cluster vs. those in all other clusters (**Fig. 4E**). In particular, Cluster 1 was distinguished by MPeM CD8^+^ T cells with high expression for checkpoint inhibitors PD-1, TIM3, and TIGIT which are characteristic of exhausted MPeM CD8^+^ T cells (34). Among the most significant differences in inferred TF activities associated with this cluster were increased values for *TCF7* (35, 36), *STAT6, BCL3, and FOS* (31-34) (FDR-corrected *p* < 10^−50^, t-test), some of which have previously been reported as TFs downstream of PD-1 (31-36) (**Fig. 4E**). Similarly, unsupervised clustering using cell surface protein expression patterns identified five major populations of MPM CD8^+^ T cells (**Fig. 3G**). Cluster 4 was distinguished by MPM CD8^+^ T cells with high expression for checkpoint inhibitors PD-1, TIM3, and TIGIT which is characteristic of exhausted MPM CD8^+^ T cells(34). Similar to MPeM CD8^+^ T cells, among the most significant differences in inferred TF activity associated with this cluster was increased *TCF7 (35, 36)* (FDR-corrected *p* < 10^−50^, t-test) (**Fig. 4G-H**).

Our analysis suggests that cell-surface receptors including those targeted by checkpoint inhibitors in tumor immunotherapy (e.g. PD-1) or T-cell co-stimulatory molecules (e.g. ICOS (inducible T-cell COStimulator (37)) can couple with common and context-dependent downstream TFs within a given cell type but in different tissue contexts (**Fig. 4C**,**F**,**I**). For example, SPaRTAN-predicted *HIF1A* activity was correlated with PD-1 protein expression in PBMC, MePM and MPM CD8^+^ T cells (38, 39). In contrast, SPaRTAN-predicted *CTCF, CREB3, CREB3L1, NR2F6, TCF7* (36), and *STAT5B* (40) activity was highly correlated with PD-1 protein expression in MPM and MPeM CD8^+^ T cells. There were also novel TFs correlated with PD-1 protein expression in CD8^+^ T cells only in one tissue type, including *FOS, MYC, RARA, CEBPB, TBP, BCL3, SREBF1, GATA6, KLF5, ZHX2, TFAP4, NR2C2, MAF* (41) and *ZNF217* for MPeM; *MAFF, THAP11* and *RFXAP* for MPM (**Supplementary Fig. 18)**.

## Discussion

SPaRTAN is a generally applicable method for exploiting parallel single-cell proteomic and transcriptomic data (based on CITE-seq) with cis-regulatory information (e.g. TF – target gene priors) to predict the coupling of TF activities with signaling receptors and pathways. Once SPaRTAN is trained using cell-type specific datasets, we can utilize the context-specific models to represent individual cells in terms of surface protein expression and TFs’ activities. These representations generate hypotheses focused on signaling regulated TF expression and function that can be testable in the laboratory or in the clinic. Application of SPaRTAN to CITE-seq datasets helps to (i) decipher critical regulators (e.g., TFs, surface receptors) underlying cellular identities (e.g. naïve vs memory T cells); (ii) determine whether given cell types have different or common regulators across tissues (e.g. B cells in spleen vs lung); (iii) determine commonalties as well as differences of cell-specific regulatory programs across healthy individuals and those manifesting a disease.

We used SPaRTAN to delineate surface receptors and TF relationships in various types of immune cells in the blood of healthy individuals. This analysis can serve as a set of reference states for such cells in settings of infectious disease or various inflammatory and autoimmune disorders. SPaRTAN was also used to analyze CITE-seq datasets generated from malignant peritoneal and pleural mesotheliomas. Malignant mesothelioma is a rare and aggressive cancer, that has not previously subjected to extensive single-cell profiling and computational analyses. Our combined analyses of immune cells in the blood and the tumor microenvironment suggests that signaling receptors e.g. CD27 and PD-1 can be coupled to common or distinct downstream TFs in different cell types.

The method we describe has several limitations. First, our analysis uses curated TF target-gene interactions (17) to determine the set of TFs that potentially regulate each gene. Those interactions were curated and collected from different types of evidence such as literature curated resources, ChIP-seq peaks, TF binding site motifs, and interactions inferred directly from gene expression. The SPaRTAN framework can be extended using scATAC-seq or bulk ATAC-seq from sorted cells for more accurate representation in both promoter and enhancer regions as performed in the context of patient-specific predictive regulatory models (42)). Our model also currently rests on the assumption that a TF either induces or represses its targets, but some TFs may play either role depending on their coordination with co-factors. Moreover, cooperatively assembling TFs (e.g., AP-1−IRF complexes(43)) can be biologically more important for fine tuning of gene expression.

SPaRTAN analysis is limited to ∼100 surface proteins (for which CITE-seq-validated barcoded antibodies are commercially available from Biolegend). However, the combination of TFs and surface proteins recovers a broad and extensive array of pathways associated with immune cell states in peripheral blood and in the tumor microenvironment. CITE-seq is currently limited to detect surface-protein and gene expression but antibodies directed against intracellular proteins will be added to future iterations of this system (44, 45) and can be easily integrated into our approach. Further, we can identify regulators by querying known pathways for upstream and downstream components of the surface protein - TF axis/connection.

Despite these limitations, SPaRTAN will accelerate the analysis of regulatory states of cells that are controlled by the reciprocal interplay of signaling systems and signal-regulated TFs. It can be used to discover new molecular connections in signal regulated gene expression programs as well as to analyze the cross-talk between signaling pathways.

## Methods

### Data and preprocessing

To construct the TF – target gene prior matrix, we downloaded a gene set resource containing TF target-gene interactions from DoRothEA (17). Those interactions were curated and collected from different types of evidence such as literature curated resources, ChIP-seq peaks, TF binding site motifs, and interactions inferred directly from gene expression. This TF – target gene prior matrix defines (**D**) a candidate set of associations between TFs and target genes. CITE-seq data for 5k (Chemistry v3), 5k (Nextgen) PBMC obtained from 10× Genomics website (**Supplementary Table 1**). A total of ∼5,000 cells from a healthy donor were stained with 29 TotalSeq-B antibodies, including CD3, CD4, CD8a, CD11b, CD14, CD15, CD16, CD19, CD20, CD25, CD27, CD28, CD34, CD45RA, CD45RO, CD56, CD62L, CD69, CD80, CD86, CD127, CD137, CD197, CD274, CD278, CD335, PD-1, HLA-DR, and TIGIT. Cell-matched scRNA-Seq data are available.

To further evaluate the validity of our method. We generated an in-house CITE-seq dataset of human malignant peritoneal and pleural mesothelioma under IRB approval from the University of Pittsburgh. Cells were stained with Totalseq-C from BioLegend and are prepared using the 10x Genomics platform with Gel Bead Kit V2 (as described below). Forty-six surface markers are measured for every cell: adt-CD274 (B7-H1, PD-L1), adt-CD273, adt-CD30, adt-CD40, adt-CD56, adt-CD19, adt-CD14, adt-CD11c, adt-CD117, adt-CD123, adt-CD194 (CCR4), adt-CD4, adt-CD25, adt-CD279, adt-TIGIT, adt-CD20, adt-CD195 (CCR5), adt-CD185 (CXCR5), adt-CD103 (Integrin αE), adt-CD69, adt-CD62L, adt-CD197, adt-CD161, adt-CD152 (CTLA-4), adt-CD223 (LAG-3), adt-CD27, adt-CD95, adt-CD134 (OX40), adt-HLA-DR, adt-CD1c, adt-CD11b, adt-CD141, adt-CD314, adt-CD66b, adt-CD366, adt-CD278, adt-CD39, adt-KLRG1, adt-CD137, adt-CD254, adt-CD357, adt-CD28, adt-CD38, adt-CD127 (IL-7Rα), adt-CD15 and adt-TCRVdelta2.

### Training cell-type specific SPaRTAN models

We trained cell-specific SPaRTAN models using affinity regression (AR) which is an algorithm for efficiently solving a regularized bilinear regression problem **(13, 16)** for any problem where the observed data can be explained as interactions between two kinds of inputs and defined here as follows. For a given set of genes and their expression profiles, we will use AR to learn an interaction matrix between cell surface proteins and TFs that explains target gene expression in cells. For a data set of *M* cells profiled using scRNA-seq across *N* genes, we let **Y ∈***R*^*N*x*M*^ be the log normalized gene expression profiles for all cells, where each column of Y corresponds to a scRNA-seq experiment. We define a matrix **D ∈***R*^*N*x*Q*^, where each row represents a gene and each column is a binary vector representing the target genes of a TF. We used curated TF target-gene interactions (17) to determine the set of TFs that potentially regulate each gene (**D**). Finally, we define a matrix P ∈*R*^*M*x*S*^ of cell surface protein attributes where each row represents a cell and each column represents protein expression levels of a surface protein across cells from scADT-seq. P, Y, and D vectors were all unit normalized. Then We set up a bilinear regression problem to learn an interaction matrix W ∈ *R*^*Q*x*S*^ for pairs of TF-surface protein features that predicts target gene expression.

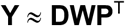

We can transform the system to an equivalent system of equations by reformulating the matrix products as Kronecker products

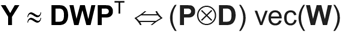

where ⊗ is a Kronecker product, and vec(.) is a vectorizing operator that stacks a matrix and produces a vector, yielding a standard (if large-scale) regression problem. To reduce the dimensionality, we reduced to a smaller system of equations where the output is the set of pairwise similarities **Y**^T^**Y** between examples rather than **Y** itself.

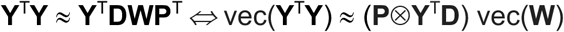

Then we used elastic-net regression to solve for the interaction matrix (**W**) and evaluate performance with 5-fold cross-validation.

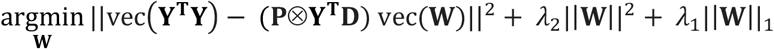

We can further reduce dimension for larger CITE-seq datasets by subjecting the feature matrix **P** and **D** to singular value decomposition prior to training. Full details and a derivation of the reduced optimization problem are provided elsewhere (16).

We can use the trained **W** to obtain different views of a CITE-seq data set: to infer the TF activities in each cell, we can right-multiply the surface protein expression profiles through the model by **WP**^T^; To infer protein activities in each cell, we can left-multiply the gene expression profile and TF target-gene interaction matrix through the model by **Y**^T^**DW** (**Fig. 1C**). We refer to these operations as “mappings” onto the TF space and the surface protein space, respectively.

### Tissue Processing of Malignant Mesothelioma

Tumor samples were washed in RPMI containing antibiotics such as amphotericin B and penicillin-streptomycin for 30 minutes followed by mechanical and enzymatic digestion and further passage via a 100µm filter. Isolated tumor infiltrating lymphocytes (TIL) were then washed in RPMI media twice, and stained with aforementioned TotalSeqC antibodies, and CD45-PE, EpCAM, and cell viability dyes. After washing, immune cells were sorted based on CD45 and utilized for sequencing library preparation.

### Simultaneous protein and transcriptomic single cell profiling of Malignant Mesotheliomas via CITE-seq

Combined surface protein and mRNA expression single cell analysis was performed using CITE-seq methodology as previously described(1). Generation of scRNAseq libraries: Live CD45^+^, EpCAM^-^ cells (i.e. all immune cells) and live EpCAM^+^ (tumor cells) were sorted from tumor tissue. Single-cell libraries were generated utilizing the chromium single-cell 5’ Reagent (V2 chemistry). Briefly, sorted cells were resuspended in PBS (0.04% BSA; Sigma) and then loaded into the 10X Controller for droplet generation, targeting recovery of 5,000 cells per sample. Cells were then lysed and reverse transcription was performed within the droplets and cDNA was isolated and amplified in bulk with 12 cycles of PCR. Amplified libraries were then size-selected utilizing SPRIselect beads, and adapters were ligated followed by sample indices. After another round of SPRIselect purification, a KAPA DNA Quantification PCR determined the concentration of libraries. The supernatant after first SPRIselect beads, containing ADTs, was used to generate ADT library.

#### Sequencing of single-cell libraries

Libraries were diluted to 2nM and pooled for sequencing by NextSeq500/550 high-output v2 kits (UPMC Genomic Center) for 150 cycles (parameters: Read 1: 26 cycles; i7 index: 8 cycles, Read 2: 98 cycles). The prepared assay is subsequently sequenced on aNextSeq500/550 with a depth of 50K reads per cell. Raw sequence data were processed via CellRanger 3.0 (10X Genomics) and aligned to GRCh38 to generate UMI matrix for the downstream analysis. Cell barcodes with fewer than 3 UMI counts in 1% of cells were removed. Antibody counts for CITE-seq were counted using CITE-seq-count. Normalization and initial explanatory analysis of CITE-seq data were performed using the Seurat R package(46). Antibody-derived tags (ADTs) for each cell will be normalized using a centered log ratio (CLR) transformation across cells. During quality control, we excluded cells with > 5000 and <300 expressed genes. Seurat “FindClusters” was applied to the first 50 principal components, with the resolution parameter set to 1. Cell labels were assigned using marker genes protein and gene expression levels.

### Significance analysis for TF activities

To assess the statistical significance of the inferred TF activities obtained from the model via the **WP**^T^ mapping, we developed an empirical null model as follows. First, we generated random permutations of the gene expression profiles Y for each cell type. For each permuted Y response matrix, we trained an AR model using true D and P input matrices and computed the corresponding inferred TF activities via the **WP**^**T**^ mapping. Using this permutation and model fitting procedure 5000 times, we generated an empirical null model for activity distribution for each cell. To identify significant TF activities, we assessed the nominal P-value for each cell relative to the empirical null model for the particular regulator TF, and we corrected for multiple hypothesis testing of non-independent hypotheses using the Bonferroni correction procedure. Then, we reported the significant regulators using an adjusted P-value of 0.15. We calculated, for each TF regulator, the frequency over samples where the regulator passed its significant threshold for a given cell type. We used this approach to identify significant TF regulators in each cell type to identify the shared and cell type-specific roles TFs.

### Flow Cytometry validation

Single cell suspensions were stained with antibodies against surface proteins (list of ab markers and clone; **Supplementary Table 3**) for 30 minutes at 4°C Dead cells were discriminated by staining with Fixable Viability Dye (eBioscience) in PBS. The cells were washed, and fixed, and permeabilized) for 1 hour followed by two wash steps with permeabilization buffer (eBioscience). Then, intracellular staining of transcription factors were conducted for 30 minutes at 4°C. Flow cytometry analysis was performed by using a Fortessa II (BD Bioscience). Flow cytometric data analyses were performed with FlowJo (Tree Star).

### Statistical analysis

Statistical tests were performed with the R statistical environment. For population comparisons of inferred TF activities and surface protein expression, we performed two-tailed Wilcoxon’s signed-rank tests and determined the direction of shifts by comparing the mean of two populations.

## Data availability

The published human PBMC CITE-Seq dataset that supports the finding of this study can be downloaded from the 10× Genomics website (https://support.10xgenomics.com/single-cell-gene-expression/datasets/3.1.0/5k_pbmc_protein_v3; https://support.10xgenomics.com/single-cell-gene-expression/datasets/3.1.0/5k_pbmc_protein_v3_nextgem). The in-house MPeM and MPM CITE-seq data is available in GEO (https://www.ncbi.nlm.nih.gov/geo/, GSExx).

## Code availability

The software for SPaRTAN is available from https://github.com/osmanbeyoglulab/SPaRTAN

## Acknowledgments

This work was supported by awards from Innovation in Cancer Informatics (ICI) Funds and R00 CA207871. We thank Dario Vignali, Tullia Bruno, Anthony Richard Cillo and Feng Shan for helpful discussions. We also thank National Mesothelioma Virtual Bank (NMVB) for supplying fresh tumor samples.

## Author contributions

H.U.O. designed the study, developed algorithmic approaches, performed all computational experiments and analyses and wrote the manuscript. X.M. implemented python version of SPaRTAN. A.S. and Z.Q. performed the experimental validation and helped to analyze results and write the experimental validation section. H. S. helped to analyze results, structure and edit the manuscript.

